# A recombinant ‘ACE2 Triple Decoy’ that traps and neutralizes SARS-CoV-2 shows enhanced affinity for highly transmissible SARS-CoV-2 variants

**DOI:** 10.1101/2021.03.09.434641

**Authors:** Shiho Tanaka, Gard Nelson, Anders Olson, Oleksandr Buzko, Wendy Higashide, Annie Shin, Marcos Gonzales, Justin Taft, Roosheel Patel, Sofija Buta, Marta Martin-Fernandez, Dusan Bogunovic, Patricia Spilman, Kayvan Niazi, Shahrooz Rabizadeh, Patrick Soon-Shiong

## Abstract

The highly-transmissible SARS-CoV-2 variants now replacing the first wave strain pose an increased threat to human health by their ability, in some instances, to escape existing humoral protection conferred by previous infection, neutralizing antibodies, and possibly vaccination. Thus, other therapeutic options are necessary. One such therapeutic option that leverages SARS-CoV-2 initiation of infection by binding of its spike receptor binding domain (S RBD) to surface-expressed host cell angiotensin-converting enzyme 2 (ACE2) is an ACE2 ‘decoy’ that would trap the virus by competitive binding and thus inhibit propagation of infection. Here, we used Molecular Dynamic (MD) simulations to predict ACE2 mutations that might increase its affinity for S RBD and screened these candidates for binding affinity *in vitro*. A double mutant ACE2(T27Y/H34A)-IgG_1_F_C_ fusion protein was found to have very high affinity for S RBD and to show greater neutralization of SARS-CoV-2 in a live virus assay as compared to wild type ACE2. We further modified the double mutant ACE2 decoy by addition of an H374N mutation to inhibit ACE2 enzymatic activity while maintaining high S RBD affinity. We then confirmed the potential efficacy of our ACE2(T27Y/H34A/H374N)-IgG_1_F_C_ Triple Decoy against S RBD expressing variant-associated E484K, K417N, N501Y, and L452R mutations and found that our ACE2 Triple Decoy not only maintains its high affinity for S RBD expressing these mutations, but shows enhanced affinity for S RBD expressing the N501Y or L452R mutations and the highest affinity for S RBD expressing both the E484K and N501Y mutations. The ACE2 Triple Decoy also demonstrates the ability to compete with wild type ACE2 in the cPass™ surrogate virus neutralization in the presence of S RBD with these mutations. Additional MD simulation of ACE2 WT and decoy interactions with S RBD WT or B.1.351 variant sequence S RBD provides insight into the enhanced affinity of the ACE2 decoy for S RBD and reveals its potential as a tool to predict affinity and inform therapeutic design. The ACE2 Triple Decoy is now undergoing continued assessment, including expression by a human adenovirus serotype 5 (hAd5) construct to facilitate delivery *in vivo*.

**Summary sentence:** An ACE2(N27Y/H34A/H374N)-IgG_1_F_C_ fusion protein decoy sustains high affinity to all SARS-CoV-2 spike receptor binding domain (RBD) protein variants tested, shows enhanced affinity for the N501Y and L452R variants, and the highest affinity for combined N501Y and E484K variants.

## INTRODUCTION

SARS-CoV-2 variants have rapidly swept the globe ^1–3^ and very recent investigations reveal that several of these variants have shown the ability to escape neutralization by convalescent antibodies in recovered COVID-19 patients ^4^ and recombinant neutralizing antibodies (nAbs) developed as therapeutics. ^5,6^ There are also fears that current vaccines may not be as effective against some of the variants and early evidence suggests that for some vaccines, this risk may exist. ^7,8^ The latter is a particular concern, as the massive vaccine efforts currently underway employ vaccines designed to elicit immune responses against first-wave sequence SARS-CoV-2 spike (S) protein and specifically the S receptor binding domain (S RBD) that binds to angiotens-inconverting enzyme 2 (ACE2) on the surface of human cells in the airway and gut that initiates viral entry and infection.^9–12^ While one response to the threat of loss of vaccine efficacy might be to continually re-design vaccines to target specific new variants, this would be an ongoing game of catch-up because it can be expected that further novel variants will emerge, particularly since several recent reports have shown that antibodies elicited by infection and vaccination act as evolutionary forces that result in the predominance of viral variants that escape these immune defenses. ^13,14^

While efforts to adapt vaccines should be encouraged, in parallel, new therapeutic approaches to neutralize viral infection that are not undermined by the presence of mutations should be advanced.

To address the need for a therapeutic and potentially prophylactic approach that has a low likelihood of being adversely affected by variant mutations, we have designed and tested ACE2 ‘decoys’ that leverage the binding of the S RBD to ACE2. This is an approach that is also being pursued by others using a variety of fusion proteins and delivery methods.^15–18^ Our ACE2 decoys under development are recombinant ACE2-IgG_1_F_C_ or -IgAF_C_ fusion proteins, with the ACE2 sequence optimized for binding affinity to S RBD. The ACE2 decoy would be given to a patient infected with SARS-CoV-2, act to prevent binding of virus to host cell ACE2 by competing with endogenous ACE2 for spike binding, and allow clearance of the virus.^19–21^

To successfully compete, an efficacious ACE2 decoy would ideally have significantly higher affinity for S RBD than endogenous, host-cell expressed ACE2. To identify ACE2 mutations with a high probability of increasing affinity, we utilized our *in silico* Molecular Dynamic (MD) simulation capabilities as described in Nelson *et al*.^22^ “*Millisecond-scale molecular dynamics simulation of spike RBD structure reveals evolutionary adaption of SARS-CoV-2 to stably bind ACE2”* wherein we reported on our identification of regions of high affinity interaction between ACE2 and S RBD based on previously reported S RBD structures. ^23,24^

Because the ACE2 decoy concept is based on interaction of ACE2 with S RBD, its binding affinity and thus efficacy may also be vulnerable to changes in the SARS-CoV-2 S RBD sequence. We therefore assessed the affinity of our ACE2 decoy, as compared to wild type (WT) ACE2, for S RBD with a variety of single or multiple mutations associated with the currently predominant variants, including the B.1.351 variant expressing E484K, K417N, and N501Y mutations, ^25^ the B.1.1.7 variant (N501Y), ^1,26^ and the Cal.20.C L452R variant. ^27^

Here, we report our findings that the combined N27Y and H34A mutations of ACE2 conferred the greatest increase in affinity for S RBD of the ACE2 variants tested. Our final ACE2 Triple Decoy also included an H374N mutation to abrogate ACE2 enzymatic activity. This ACE2 Triple Decoy not only maintained affinity for variant S RBD, it showed an increased affinity for S RBD expressing N501Y or L452R mutations.

## RESULTS

### Wild type (WT) ACE2-IgG_1_F_C_ and ACE2-IgAF_C_ decoys show high affinity for the spike receptor binding domain

In initial studies to design an ACE2 decoy, we determined the affinity of both recombinant wild type (WT) ACE2(WT)-IgG_1_F_C_ and -IgAFc fusion proteins for binding to S RBD by Biolayer Interferometry (BLI) analysis. The ACE2(WT)-IgG_1_F_C_ decoy (Fig. 1A) showed high affinity for S RBD in both 1:1 binding with a coefficient of dissociation (K_D_) of 21.40 nM and binding with avidity with a K_D_ of 0.762 nM (Fig. 1C and D, respectively; values in Fig. 1F). The ACE2(WT)-IgAF_C_ dimeric fusion protein (Fig. 1B) demonstrated even higher binding (with avidity) affinity for S RBD with a K_D_ of 0.166 nM (Fig. 1E and F).

**Fig. 1.**
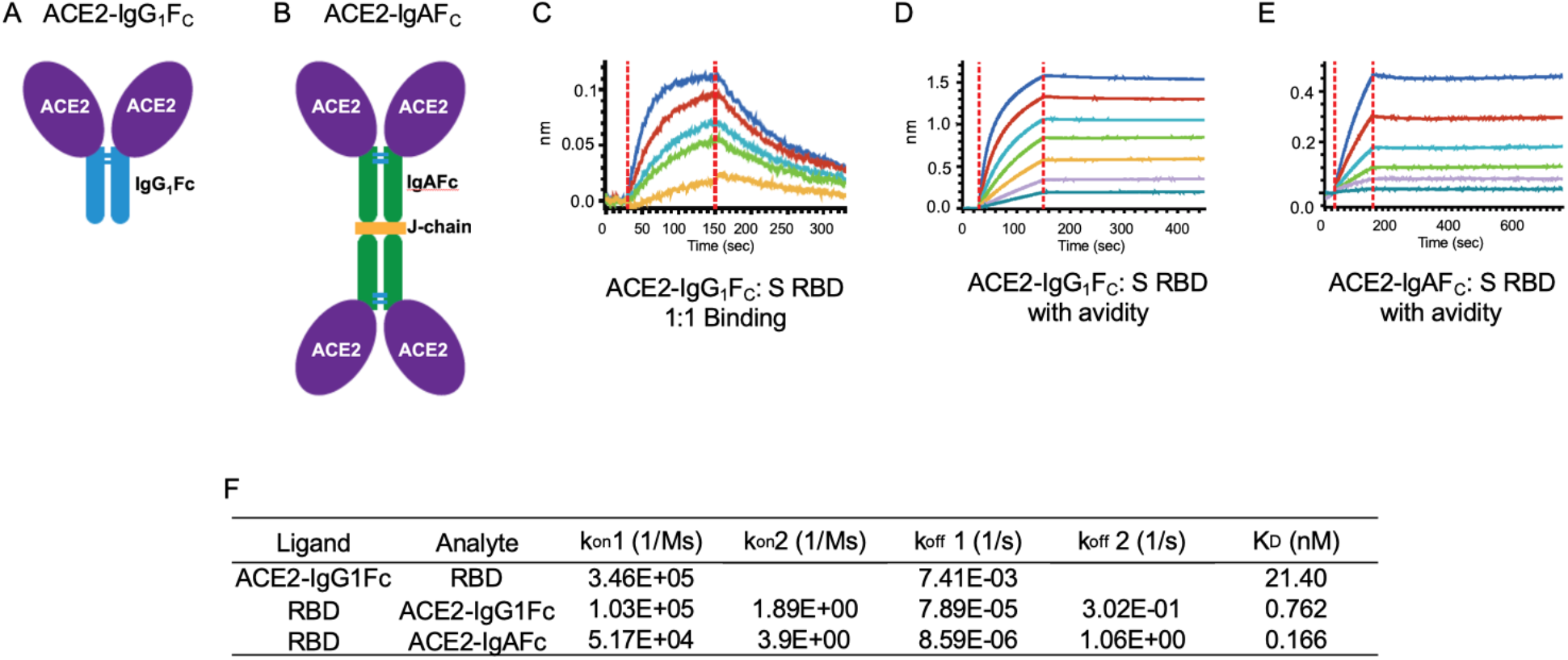
ACE2-IgG_1_F_C_ and dimeric -IgAF_C_ decoys bind the spike receptor binding domain (S RBD) with high affinity. The (A) ACE2-IgG_1_F_C_ decoy; (B) dimeric ACE2-IgAF_C_ decoy fused via a J-chain are shown. Biolayer Interferometry (BLI) kinetics analysis of (C) 1:1 binding and (D) binding with avidity for the ACE2-IgG_1_F_C_ decoy; and (E) BLI binding with avidity for the ACE2-IgAF_C_ decoy are shown. (F) Table of binding affinity values.

### An ACE2 decoy expressing T27Y and H34A mutations confers the greatest enhancement of affinity for S RBD and improved neutralization of live SARS-CoV-2 virus *in vitro*

Based on MD simulation-based predictions of mutations that may confer enhanced binding affinity of ACE2 for S RBD, several ACE2 variants were tested for binding affinity as ACE2-IgG_1_F_C_ fusion proteins. As shown in Figure 2, a tyrosine (Y) substitution for threonine (T) at residue 27 and an alanine (A) substitution for histidine (H) at residue 34 of ACE2 resulted in 3~5 fold increases in binding affinities (T27Y K_D_ = 8.01; H34A K_D_ = 4.09 nM). Combination of the T27Y and H34A substitutions results in a synergistic enhancement of binding affinity, showing an ~35-fold increase in binding affinity as compared to ACE2(WT) with the K_D_ decreasing to 0.56 nM (Fig. 2D and F).

**Fig. 2.**
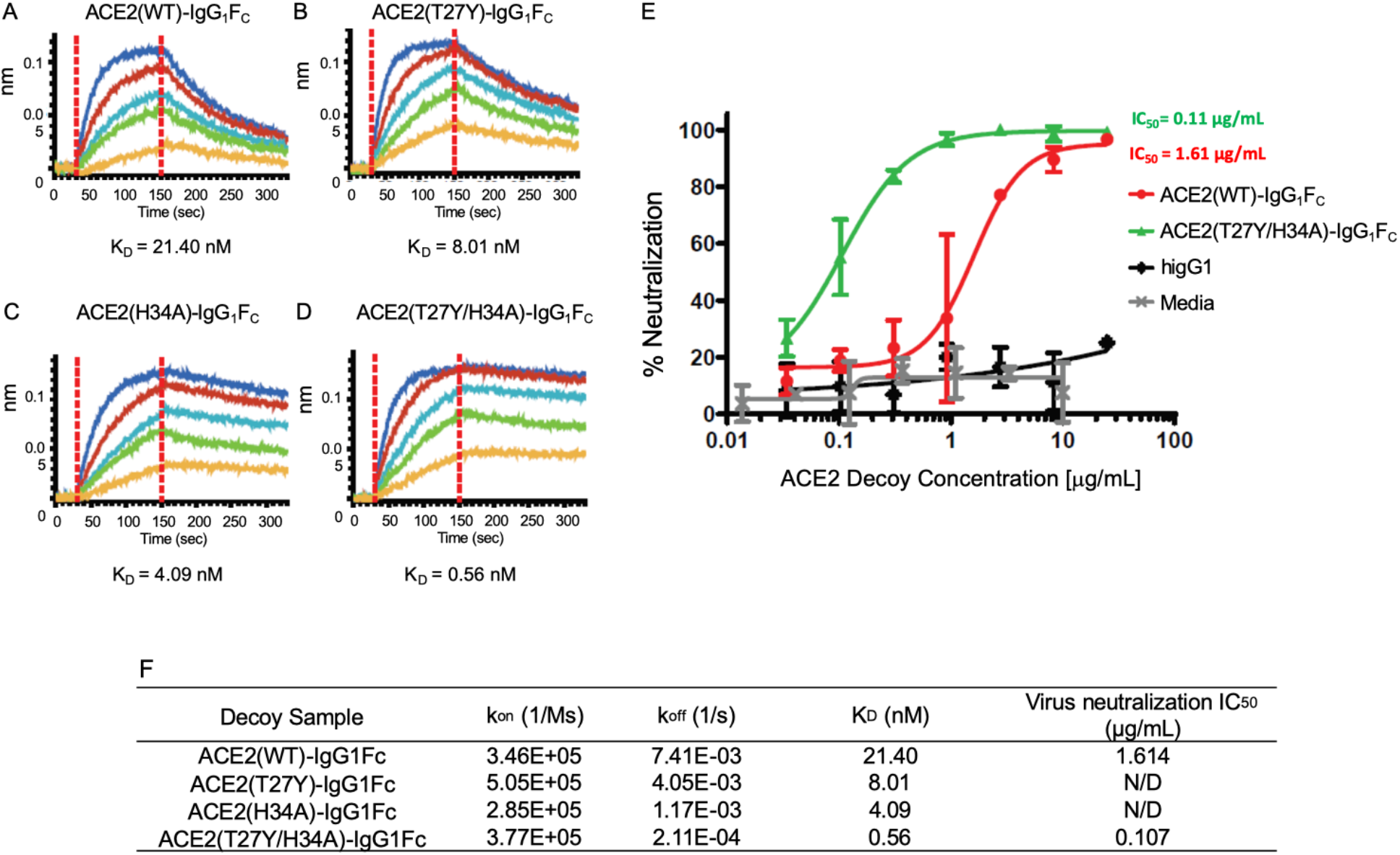
Biolayer Interferometry (BLI) of mutated ACE2-IgG_1_F_C_ decoys and the live virus neutralization assay. The kinetics of binding are shown for (A) ACE2(WT)-IgG_1_F_C_, (B) ACE2(T27Y)-IgG_1_F_C_, (C) ACE2(H34A)-IgG_1_F_C_, and (D) ACE2(T27Y/H34A)-IgG_1_F_C_ decoys. (E) The percent neutralization over increasing concentrations (μg/mL) of decoy is shown. (F) Binding affinity values. higG1 – a human IgG1 control.

The ACE2(T27Y/H34A)-IgG_1_F_C_ double decoy was compared to an ACE2(WT)-IgG_1_F_C_ decoy in a live SARS-CoV-2 virus assay using Vero E6 cells. As shown in Figure 2E, the double mutant ACE2 decoy showed ~15-fold improvement in SARS-CoV-2 neutralization capability compared to the wild type ACE2 Decoy.

### MD simulations provide insight into greater affinity of T27Y and H34A ACE2 for S RBD

MD simulations (Fig. 3) of the ACE2 T27Y and H34A substitutions suggest that a tyrosine (Y) substitution for threonine (T) at residue 27 introduces favorable hydrophobic contacts with RBD. An alanine (A) substitution for histidine (H) at residue 34 of ACE2 allows more surface area for RBD residues to contact the ACE2 helix and may favorably increase entropy by increasing side chain flexibility. Synergy between these mutations occurs since their effects are independent and do not perturb the binding pose.

**Fig. 3.**
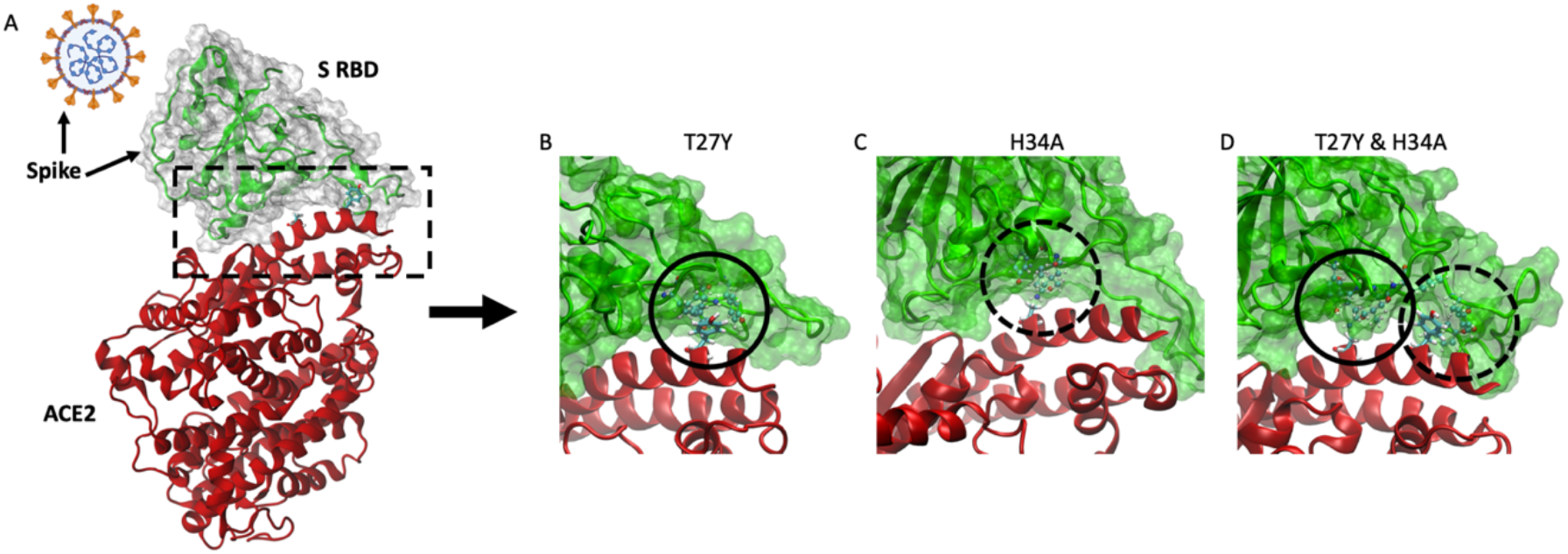
Molecular effects of T27Y and H34A ACE2 mutations predicted by MD simulation. (A) Spike (S) occurs as a trimer on the viral surface (orange projections), with the receptor binding domain (RBD) being on the outermost surface. The interface between S RBD and ACE2 is within the dashed box. Simulation models are shown for (B) ACE2(T27Y)- (circle), (C) ACE2(H34A)- (dashed circle), and (D) ACE2(T27Y/H34A)-S RBD interactions.

### Addition of an H374N mutation inhibits ACE2 enzyme activity

In addition to enhanced affinity for competitive binding of S RBD, we wanted to inhibit the enzymatic activity of ACE2.^28^ Angiotensin-converting enzyme 2 has an important role in homeostasis of the renin-angiotensin system ^29–31^ by cleavage of its substrate angiotensin 1-9 ^32^ and its activity affects a variety of systems. Addition of enzymatically active recombinant ACE2 to the system presents a high risk of unwanted side effects and since S RBD binding, but not substrate cleavage activity, is the key function for the ACE2 decoy, we tested a variety of mutations predicted to inhibit ACE2 enzymatic activity with a low likelihood of affecting S RBD binding affinity.

All of the ACE2 mutations (R273Q, R273K, R273L, H345A, H505L, H374N, or H378N) predicted or known to inhibit ACE2 enzymatic activity ^33,34^ did inhibit this activity in the assay (Supplementary Methods, Supplementary Fig. S1). ACE2 triple mutant decoys comprising the S RBD binding affinity-enhancing T27Y/H34A mutations and the enzymatic activity-inhibiting mutations were produced and binding affinity assessed. Of the triple mutants, those with either the R273K or H374N mutations showed the highest S RBD affinity (Supplementary Table S1).

The final ACE2 Triple Decoy chosen for further testing was ACE2 (T27Y/H34A/H374N)-IgG_1_F_C_ due to its more favorably biophysical characteristics as compared to an R273K-containing triple mutant, including a lower propensity to aggregate and a higher titer (Supplementary Fig. S2 and Table S2).

### The ACE2 Triple Decoy shows enhanced binding to S RBD expressing N501Y and L452R variants, with the highest affinity for S RBD expressing both N501Y and E484K

The binding affinities of both the wild type ACE2 decoy (ACE2(WT)-IgG_1_F_C_) and the ACE2(T27Y/H34A/H374N)-IgG_1_F_C_ Triple Decoy to S RBD WT or a series of mutations found in the B.1.351/P.1 ^35^ (E484K/K417N/N501Y), B.1.1.7 (N501Y), ^1,26^ and CAL.20.C (L452R) ^27^ variants are shown in Figure 4.

**Fig. 4.**
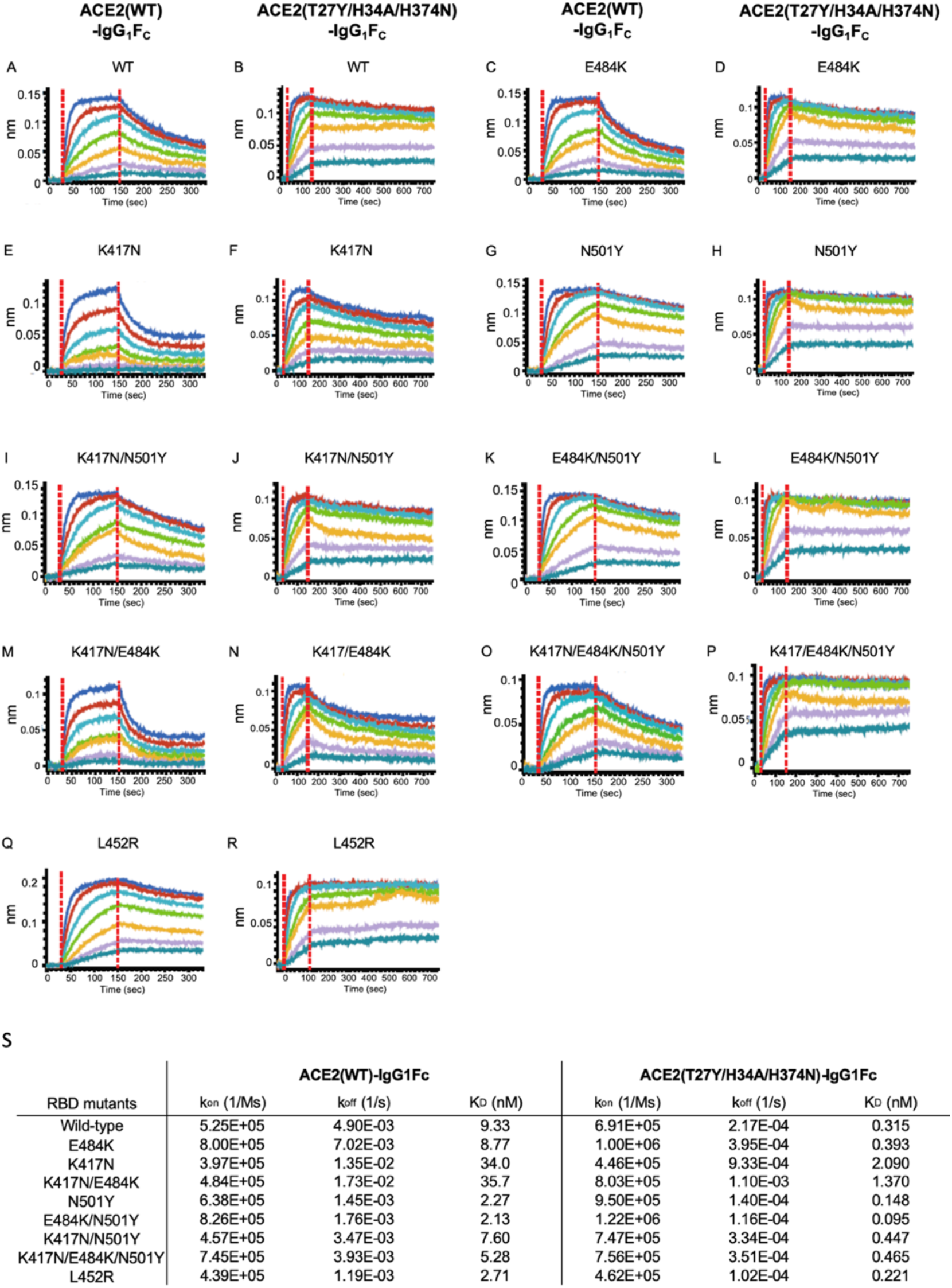
Biolayer Interferometry (BLI) analysis of ACE2(WT)- and ACE2(T27Y/H34A/H374N)- IgG_1_F_C_ to spike receptor binding domain (S RBD) variants. (A-R) Comparative binding by the ACE2(WT)-IgG_1_F_C_ or the ACE2(T27Y/H34A/H374N)-IgG_1_F_C_ Triple Decoy to S RBD WT or a series of mutations (E484K, K417N, N501Y) alone and in combination; or S RBD with the L452R mutation are shown side-by-side (for example, ACE2(WT)IgG_1_F_C_ versus ACE2(T27Y/H34A/H374N)-IgG_1_F_C_ binding to S RBD WT are shown in A and B). (S) Binding affinity values.

The ACE2 Triple Decoy showed higher binding affinity to all S RBD sequences as compared to the wild type ACE2 decoy. As compared to the ACE2 Triple Decoy binding affinity for S RBD WT, affinities for S RBD E484K/N501Y, N501Y alone and L452R were higher; affinities for S RBD E484K, K417N/N501Y, N417N/E484K/N501Y, K417K/E484K, and K417N were lower. Findings were similar with the wild type ACE2 decoy, with the highest affinity seen for E484K/N501Y and N501Y alone, and the lowest affinities for variants expressing K417N. N501Y and L452R showed ~2-3 fold increase in binding affinity for both wild type ACE2 decoy and ACE2 Triple Decoy. E484K alone did not affect binding affinity to ACE2. K417N weakened binding affinity for ACE2 (WT) and triple decoys, but affinity was restored when combined with N501Y. The E484K, K417N and N501Y mutations occur together in the B.1.351 strain, whereas L452R alone is found in CAL.20.C, therefore assessment of ACE2 WT binding to these variants as they occur in nature is most physiologically relevant (Supplementary Fig. S3 and Table S3).

### Inhibition of ACE2:S RBD binding in the surrogate virus neutralization assay correlates with binding affinity

The surrogate SARS-CoV-2 neutralization assay cPass™ ^36^ is based upon assessment of inhibition of binding of ACE2 (WT) to A SRB (WT). It is typically used to ascertain the presence of anti-S RBD antibodies in serum. Such antibodies inhibit binding of S RBD to ACE2 bound to an ELISA plate, and inhibition of greater than 20% has been reported to correlate with neutralization of live virus. Here, the surrogate assay was used to determine if the ACE2 Triple Decoy could inhibit S RBD WT and variant binding to plate-bound ACE2, that is, compete with ACE2 (WT) for S RBD binding.

As shown in Figure 5, the ACE2 Triple Decoy inhibition of binding by the ACE2 Triple Decoy was similar for S RBD WT, E484K, and L452R; and only slightly lower for S RBD N501Y and E484K/K471N/N501Y. Only S RBD K471N binding showed a lower level of inhibition by the ACE2 Triple Decoy, all other variants tested showed inhibition that was significantly higher than the no-decoy control.

**Fig. 5.**
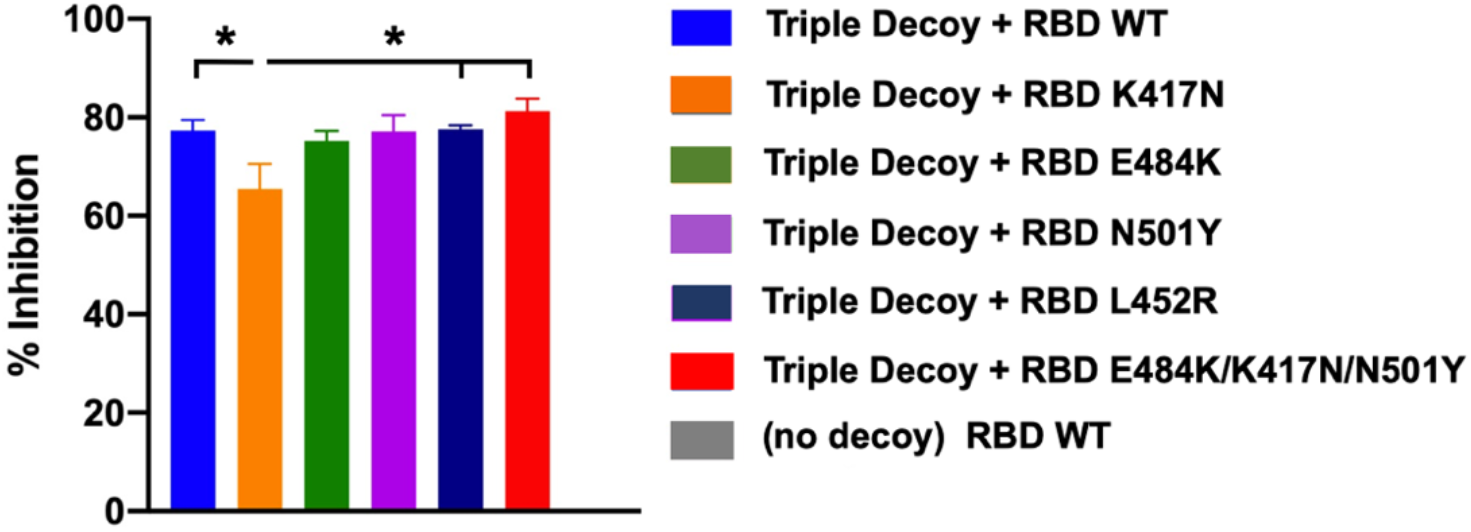
Inhibition of spike receptor binding domain (RBD) wild type (WT) and RBD variant binding to ACE2 by the ACE2 Triple Decoy in the surrogate neutralization assay. The percent inhibition of (competition for) RBD binding to ACE2 bound to the ELISA plate in the surrogate virus neutralization assay cPass™ is shown for the ACE2 Triple Decoy with S RBD WT and the listed variants. All RBD concentrations were 25 μg/mL. The negative control has no decoy. Statistics performed using One-way ANOVA and Tukey’s post-hoc analysis to compare Triple Decoy (but not ‘no decoy’) binding to RBD WT and variants. For RBD K417N vs WT, p = 0.0495; vs L452R, p = 0.0451; and vs E484K/K417N/N501Y, p = 0.0128.

### MD simulation accurately predicts the relative affinities confirmed by *in vitro* testing

We used Adaptively-Biased MD (ABMD) simulations, ^37^ which allow observation and quantification of binding and unbinding, of both ACE2 WT and ACE2 (T27Y/H34A) binding to S RBD WT or B.1.351 to predict binding affinities. For these simulations, the B.1.351 variant comprising the E484K, K417N, and N501Y mutations was used because these mutations occur together naturally and thus this combination has high physiological relevance. The ACE2 T27Y/H34A sequence without the additional H374N enzyme-deactivating mutation found in the ACE2 Triple Decoy was used because earlier simulations had been unable to detect a change in affinity due to the presence of the H374N mutation.

We used the Helmholtz binding free energy (ΔA_bind_), determined by the ratio of the probability of the bound and unbound states based on the Free Energy Surfaces (FES) (Figure 6), to assess relative predicted affinities, where more negative values of ΔA_bind_ indicate a stronger association. Details of the ABMD simulations and Helmholtz calculation can be found in *Methods*. The calculated free energies of binding, in order of predicted affinity from lowest to highest, are: ACE2 WT:RBD WT (−4.1 kcal/mol; Fig. 6A); ACE2 WT:RBD B.1.351 (−5.1 kcal/mol; Fig. 6B); ACE2 T27Y/H34A:RBD B.1.351 (−6.3 kcal/mol; Fig. 6C); and ACE2 T27Y/H34A:RBD WT (−7.0 kcal/mol; Fig. 6D).

**Fig. 6.**
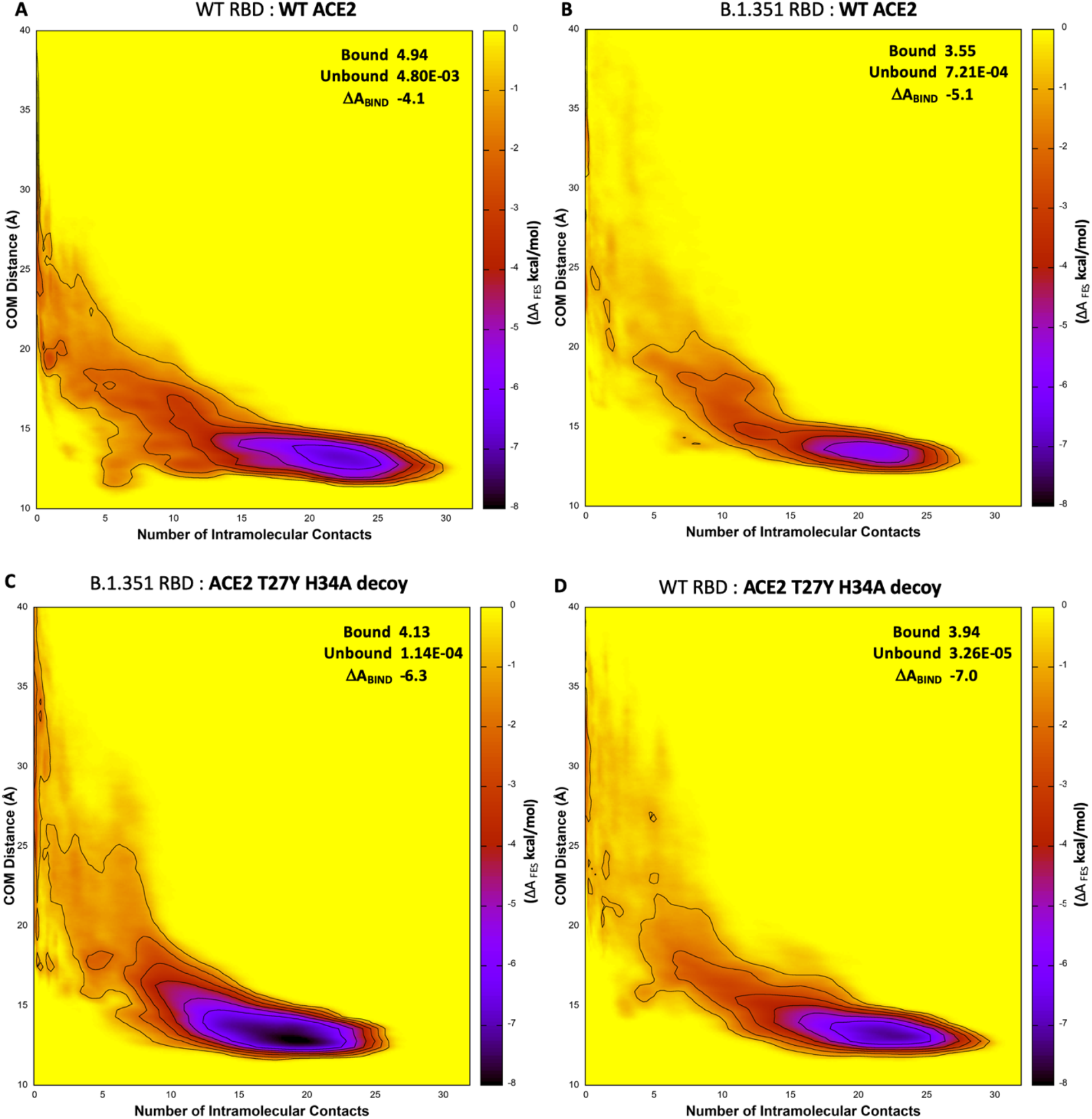
MD simulation predicts highest affinity for the T27Y/H34A decoy to S RBD WT and B.1.351. The free energy surfaces (FES) of wild type (WT) ACE2 upon interaction with (A) WT RBD or (B) B.1.351 RBD; and FES for the ACE2 T27Y/H34A decoy and (C) B.1.351 RBD or (D) WT RBD are shown. Darker purple represents lower free energy (ΔA_FES_, scale at right of each panel). The free energy is a function of the number of intramolecular contacts (x-axis) and the distance between the centers of mass (COM, y-axis) of the interface regions.

The predictive utility of these simulations is supported by the findings from the affinities (KD) determined *in vitro* and presented in Figure 4, where (for the combinations tested in MD simulations) the lowest affinity was also seen for ACE2 WT : RBD WT (K_D_ = 9.33 nM), followed by ACE2 WT : RBD B.1.351 (K_D_ = 5.28 nM), then ACE2 Decoy : RBD B.1.351 (KD = 0.465 nM), and ACE2 Decoy : RBD WT (KD = 0.315 nM). Note that all affinities were high, and higher for Triple Decoy binding than ACE2 WT for all RBD sequences tested.

## DISCUSSION

To our knowledge, we are the first to report binding affinities of a recombinant mutant ACE2 decoy to the spike receptor binding domain expressing N501Y, E484K, N417Y, or L452R mutations; although we note Huang *et al*. reported previously on the affinity of their ACE2-FC to S RBD with the D614G mutation. ^38^ The greater affinity of ACE2 for S RBD with the N501Y substitution alone or in combination with E484K reported here is in alignment with our findings in Nelson *et al*,^39^ wherein we used MD simulation to predict that these mutations have a high probability of increasing affinity for ACE2.

The MD simulation data presented here used to guide design of the ACE2 Triple Decoy and to predict affinities of the decoy as compared to ACE2 WT for a series of variants reveal again the merits of such simulations as a tool to inform therapeutic design.

Interestingly, widespread use of an ACE2 decoy has the potential itself to act as an evolutionary force; however, an ACE2 decoy largely recognizes the same residues as endogenous ACE2 and therefore it is highly unlikely a SARS-CoV-2 variant could emerge that ‘escapes’ the decoy yet still binds to endogenous ACE2. This phenomenon along with limited use of a decoy for therapy as compared to the spread of virus in a large population with opportunity for selection, makes the decoy approach less vulnerable to loss of efficacy due to mutation of the virus.

The enhanced binding affinity of our Triple Mutant ACE2 Decoy to S RBD with the variant mutations tested here supports continued pursuit of this therapeutic approach and further provides hope that even should the efficacy of vaccines currently in distribution or therapeutic neutralizing antibodies raised against WT spike be lessened by these variants, there will be an alternative therapeutic approach to successfully treat COVID-19 disease.

In our next steps in development of the ACE2 Triple Decoy, we will address the challenge of stability and successful delivery. Others developing ACE2 decoys have suggested use of intranasal ^40^ or nanoparticle/extracellular vesicle delivery. ^16,41–43^ We anticipate going forward into our next studies using the dimeric IgA ^44^ fusion protein decoy expressed by the human adenovirus serotype 5 E1, E2b, E3 deleted (hAd5 [E1-, E2b-, E3-]) platform that we have used successfully in our vaccine development. ^45,46^ This platform can readily be used to generate oral and/or intranasal formulations to further facilitate delivery. Our ACE2 Triple Decoy delivered *in vivo* using the hAd5 platform is anticipated to overcome barriers to successful delivery and will be tested in animal models of SARS-CoV-2 infection in future studies.

## METHODS

### MD simulation

#### System Setup

The WT-ACE2/RBD complex was built from the cryo-EM structure, PDB 6M17 of full-length human ACE2 in the presence of the neutral amino acid transported B^0^AT1 with the S RBD as shown in Yan *et al*. ^47^ using RBD residues 336-518 and ACE2 residues 21-614. ACE2 residues 27 and 34 were mutated to tyrosine and alanine, respectively. The final simulation system was built using the Amber ff14SB force field ^48^. The RISM program from AmberTools19 ^49^ was used to determine optimal locations for water molecules in direct contact with the proteins. Bulk waters were added to create a sufficient octahedral water box and sodium ions were added at random locations to neutralize the system. After introducing mutations at the relevant residues, the same procedure was used to generate the other three systems.

#### Simulation

Ten copies of each RBD/ACE2 complex were minimized, equilibrated and simulated. Minimization occurred in two phases. During the first, the protein and RISM-placed waters were restrained. The second phase minimized the entire system. Dynamics then began and the temperature was ramped from 0 to 300K while restraining the protein and RISM-placed waters. All dynamics used SHAKE restraints on hydrogen-containing bonds and a 2 fs timestep. All restraints were then released and the system was equilibrated in the NPT ensemble for 2 ns. Finally, each system was equilibrated in the NVT ensemble for 100 ns.

Steered MD was used to prepare the equilibrated systems for free energy calculation. Contacting residues from the adaptively biased MD (ABMD) simulations in Nelson *et al*. ^22,37^ were used. Starting from the NVT equilibrated structures and over a 10 ns simulation, the number of intermolecular contacts was linearly reduced to 0 using a 10 kcal/mol*Å steering bias. Structures were randomly selected from the steered MD simulations and used to seed ABMD simulations. Two dimensional ABMD simulations used intermolecular contacts and the center of mass distance as collective variables. Centers of mass were defined as the alpha carbons from all interfacial residues in each molecule. The well-tempered ABMD bias potential ^50^ was used for free energy calculations. ABMD simulations were run for a total of 15.6μs, 16.0μs, 16.0μs and 16.38μs for the ACE2 WT:RBD WT, RBD WT:ACE2 T27Y/H34A, RBD B.1.351:ACE2 WT and RBD B.1.351:ACE2 T27Y/H34A, respectively. Production simulations were run in the NVT ensemble meaning the calculated free energy corresponds to the Helmholz free energy. (ΔA)

ABMD produces a free energy surface (FES) that describes the relative free energy between any two points on the FES, ΔA_FES_. The binding free energy (ΔA_bind_) is determined by the ratio of the probability of the bound and unbound states and can be determined from the FES:

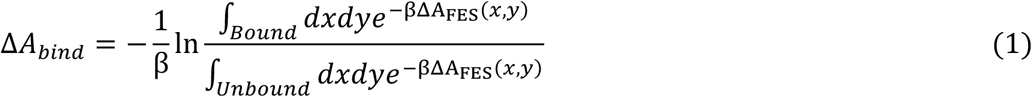

Where β is the inverse of the Boltzmann constant multiplied by the temperature in Kelvins. More negative values of ΔA_bind_ indicate a stronger association. The calculated ΔA_bind_ values can be directly compared.

The “Bound” integral in equation 1 is defined to be over all ΔA_FES_(x,y) values with the number of contacts greater than 0.05 while the “Unbound” integral is over all values with fewer than 0.05 contacts. ΔA_bind_ was calculated with different boundaries ranging from 0.0 to 1.0, inclusive. As expected, the resulting values of ΔA_bind_ changed based on the chosen boundary. However, the relative ordering of the values did not. The value of 0.05 contacts was chosen as the boundary because it allowed for unambiguous categorization of points as either “unbound” (x = 0) or “partially” or “fully bound”. All simulations were performed with the GPU-enabled version of pmemd from Amber20^49^. Multiple-walker ABMD simulations ^51^ used the MPI version of pmemd.cuda from Amber20.

### Production of ACE2 Decoys and S RBD

#### Expression constructs

Polymerase Chain Reactions (PCR) were conducted using PrimeSTAR GXL DNA Polymerase (Takara Bio) per manufacturer’s instructions. Primers and Gene Fragments were synthesized by Integrated DNA Technologies (IDT). For Gibson Assembly, NEBuilder Hifi DNA Assembly Master Mix (New England Biolabs) was used. For DNA ligation, we used T4 DNA Ligase (NEB) per the manufacturer’s instructions. Plasmid sequences were confirmed by sanger sequencing (Genewiz).

ACE2-IgG_1_F_C_ was created by Gibson Assembly of 3 fragments: 1) the vector backbone from a NheI-XhoI 7.168 kb fragment of pWT35, 2) ACE2 from a 1.86 kb PCR product of WH1043 and WH1044 amplification of gene-synthesized ACE2 codon optimized for expression in CHO epithelial cell line (AO615ACE2), and 3) IgG_1_F_C_ from a 0.701 kb PCR product of pXL159, using primers WH1045 and WH1046. ACE2 R273Q-IgG_1_F_C_ was constructed similarly, with the exception that ACE2 R273Q was created by splice by overlap extension (SOE). A 1.86 kb SOE product was created by amplification with primers WH1043 and WH1044 of two PCR products: 1) 860 bp amplification of AO615ACE2 with primers WH1043 and WH1049, and 2) 1.059 bp amplification product of AO615ACE2 with primers WH1050 and WH1044.

ACE2 T27Y/H34A-IgG_1_F_C_ was constructed by the Gibson Assembly of: 1) a 9.041 kb NheI-PshA1 digestion fragment of ACE2-IgG_1_F_C_ plasmid, and 2) a 0.773 kb SOE product of primers 5MutF and 5MutR of two PCR products. The first PCR is a 0.154 kb amplification of plasmid SR9 with primers 5MutF and ACE2T27YR). The second PCR products is a 0.642 kb amplification of plasmid SR9 with primers ACE2T27YF and 5MutR.

Most of the triple mutants were created by Gibson Assembly of three fragments: 1) the vector backbone from a 7.168 kb NheI-XhoI fragment of pWT35, 2) IgG_1_F_C_ from a 0.701 kb PCR amplification of pXL159 with primers WH1045 and WH1046, and 3) the ACE2 variant from a 1.86 bp PCR containing the three mutations. F or the latter, the mutants were amplified with primers WH1043 and WH1044 with templates pWH230 (for T27Y/H34A/R273K), pWH231 (T27Y/H34A/R273L), pWH236 (T27Y/H34A/H345A), pWH233 (T27Y/H34A/H505L), pWH234 (T27Y/H34A/H374N), and pWH235 (T27Y/H34A/H378N).

ACE2 T27Y/H34A/R273Q was constructed by ligating the 9.041 bp NheI-PshA1 fragment of ACE2 R273Q-IgG_1_F_C_ and the 0.661 kb NheI-PshA1 fragment of ACE2 T27Y/H34A-IgG_1_F_C_. *Primers* (*5’ → 3’*):

5MutF GTCTTTTCTGCAGTCACCGTCACCGTCCTTG
5MutR TGCGTGAAGATGCTCATAGAGTGGTTTT
ACE2T27YF CGAGGAGCAGGCTAAATACTTTCTGGATAAGTTTAACC
ACE2T27YR GGTTAAACTTATCCAGAAAGTATTTAGCCTGCTCCTCG
WH1043 CCGTCCTTGACACGAAGCTGCTAGCGCCACCATGAGCAGCAGTAGTTGGCT
WH1044 GGTGGGCAAGTATGTGTTTTGTCTGCATAGGGAGACCAGTCTG
WH1045 AAAACACATACTTGCCCACCTTGTCCTG
WH1046 AGTTCTAGAATCGGTATCGCTCATTTGCCAGGGCTCAGTGACAGACTC
WH1049 TGGTCCAGAACTGTCCCCACATG
WH1050 CATGTGGGGACAGTTCTGGACCA

#### Maxcyte® transient transfection

For transient expression of ACE2 decoys by Maxcyte® transfection, CHO-S cells were cultured in suspension in CD-CHO media supplemented with 8 mM L-glutamine in shaker flasks at 37 °C with 125 rpm rotation and 8 % CO_2_. For transfection, cells in the exponential growth stage were pelleted by centrifugation at 1,400 rpm for 10 min, re-suspended in 10 mL of electroporation buffer, and re-pelleted at 1,400 rpm for 5 min. The cell pellet was resuspended at a density of 2 x 10^8^ cells/mL in electroporation buffer, mixed with the plasmid harboring either the ACE2(WT)-IgG1Fc or ACE2(WT)-IgA sequence at a concentration of 150 μg/mL, and transfected using OC-400 processing assemblies in a Maxcyte® ExPERT ATx Transfection System. Transfected cells were incubated for 30 min at 37 °C, 5% CO_2_ and then resuspended in Efficient Feed A Cocktail (CHO-CD EfficientFeed™ A + 0.2% Pluronic F-68 + 1% HT Supplement + 1% L-glutamine) at a density of ~4-6 x 10^6^ cells/mL. This cell culture was incubated at 37 °C with 5% CO_2_ and 125 rpm rotation overnight, 1 mM sodium buryrate was added, and the culture was further incubated at 32 °C with 3% CO_2_ and 125 rpm for 13 more days; during this incubation period, Maxcyte® Feed Cocktail (13.9% CD Hydrolysate, 69.5% CHO CD EfficientFeed™ A, 6.2% Glucose, 6.9% FunctionMax™ Titer Enhancer, 3.5% L-Glutamine) was added at 10% of the culture volume on Days 3 and Day 8.

#### FectoPRO® transient transfection of ACE2 Mutant Decoys

For transient expression of ACE2 mutant decoys by FectoPRO® transfection, CHO-S cells in suspension were cultured in CD-CHO media supplemented with 8 mM L-glutamine in shaker flasks at 37 °C with 125 rpm rotation and 8 % CO_2_. One day before transfection, CHO-S cells were seeded at a density of 1x 10^6^ cells/mL in 45 mL culture flask. On the day of transfection, 75 μL of FectoPRO® transfection reagent (PolyPlus-transfection®) was mixed with 5 mL of 15 μg/mL pcDNA3 plasmid DNA in CD-CHO media and incubated for 10 min at room temperature. The DNA/transfection reagent mixture was added to 45 mL of CHO-S culture and incubated at 37 °C with 5% CO_2_ and 125 rpm rotation. On Day 3, 50 mL of the CD-CHO media supplemented with 8 mM L-glutamine was added and the culture incubated for an additional 4 days. *Lipofectamine® transient transfection of RBD constructs*

For transient expression of RBD wild-type and RBD mutants, HEK-293T cells were cultured and incubated at 37°C with 5% CO_2_. Plasmids harboring RBD constructs were mixed with lipofectamine with 1:1 (v:v) and incubated for 20 min at room temperature. The mixture was then added to cultures and incubated for 3-4 days.

#### Purification of ACE2 Decoy IgGs

The MaxCyte® or FectoPRO® transfection cell culture medium was centrifuged and filtered through a 0.22 μm filter to remove cells and debris, then loaded onto a HiTrap™ MabSelect SuRe™ column on the AKTA Pure system pre-equilibrated with 10 mM Na Phosphate and 150 mM NaCl at pH 7.0. After loading, the column was washed with 10 column volumes of the same buffer. The protein was eluted with 100 mM sodium acetate, pH 3.6, then immediately neutralized using 2 M Tris pH 8.0. The elution fractions were pooled and dialyzed into 10 mM Hepes and 150 mM sodium chloride at pH 7.4.

#### Purification of ACE2 Decoy IgAs

The MaxCyte® transfection cell culture medium was centrifuged and filtered through a 0.22 μm filter to remove cells and debris, then loaded to a gravity column packed with CaptureSelect® IgA resins (Thermo Fisher) pre-equilibrated with 10 mM Na Phosphate and 150 mM NaCl at pH 7.0. After loading, the column was washed with 10 column volumes of the same buffer. The protein was eluted with 100 mM sodium acetate, pH 3.0, then immediately neutralized using 2 M Tris, pH 8.0. The elution fractions were pooled and dialyzed into 10 mM Hepes and 150 mM sodium chloride, pH 7.4.

#### Purification of RBD and RBD mutants

The Lipofectamine transfection cell culture medium was centrifuged and filtered through a 0.22 μm filter to remove cells and debris. A buffer of 50 mM Tris, 100 mM sodium chloride, and 10 mM imidazole was added to the supernatant then loaded to a gravity column packed with Ni-NTA resins (Qiagen) pre-equilibrated with 20 mM Tris, 300 mM sodium chloride, and 10 mM imidazole, pH8.0. After loading, the column was washed with 10 column volumes of the same buffer. The protein was eluted with 20 mM Tris, 150 mM sodium chloride, and 300 mM imidazole. The elution fractions were pooled and dialyzed into 10 mM HEPES and 150 mM sodium chloride, pH 7.4.

### RBD affinity determination of ACE2 decoys by Bio-Layer Interferometry (BLI)

The running buffer in all experiments was 10 mM HEPES, 150 mM NaCl, pH 7.4, with 0.02% tween 20, and 0.1% BSA unless otherwise indicated. For the determination of 1:1 binding affinity of ACE2 Decoys against SARS-CoV2 RBD wild-type and mutants, ACE2 Decoys were immobilized on an AHC sensor (Sartorius Corporation) and an RBD concentration series of 200, 100, 50, 25, 12.5, 6.25, 3.125 nM was used to determine the dissociation coefficient (K_D_). For determining ACE2 Decoy binding affinity with avidity, biotinylated RBD was immobilized on streptavidin (SA) or high-precision SA (SAX) sensors, and the ACE2 Decoy concentration series of 200, 100, 50, 25, 12.5, 6.25, 3.125 nM was used to determine K_D_.

### cPass™ ^36^ surrogate SARS-CoV-2 neutralization assay

High BIND 96-well ELISA plates (Corning #3369) were coated with 50 ng/well ACE2 wild type decoy overnight at 4°C. After the antigen solution was removed, each well was blocked with 150 μL of 5% BSA/PBS for 1-2 hours at room temperature with shaking. During the blocking step, 40 μL of 50 nM RBD and RBD variants were mixed with 40 μL of 25 μg/mL of ACE2 decoy were mixed in a 96-well plate and incubated at room temperature for 30 min with shaking. After blocking, the plate was then washed 3 times with 250 μL of PBS with 0.05% Tween 20 (PBS-T). To each well, 30 μL of 1:1667 diluted mouse anti-His, HRP and 60 μL of RBD/ACE2 decoy (or a no decoy control) were added and incubated at room temperature for 30 min. The plated was washed once with 250 μL of PBS-T. To develop the signal, 50 μL of TMB solution was added and incubated at room temperature in dark for 30 min, followed by addition of 50 μL of 2M sulfuric acid; absorbance was the read at 450nm. The percent inhibition was calculated using (1-A450 (RBD+Decoy) /A450 (RBD only))x100.

## SUPPLEMENTARY MATERIAL

### Supplementary Methods

#### Assay for ACE2 enzymatic activity

Enzymatic activity ACE2 decoys expressing a variety of mutations - R273Q, R273K, R273L, H245A, H505L, H374N, and H378N – selected to inhibit activity in combination with the S RBD affinity-enhancing mutations T27Y and H34A were assessed in the FRET based ACE2 activity assay.

### Supplementary Results

As shown in Figure S1, wild type (WT) and the T27Y/H34A mutations had similar ACE2 enzymatic activity. Addition of R273Q, R273K, R273L, H245A, H505L, H374N, or H378N mutations in combination with the T27Y/H34A mutations inhibited activity of ACE2.

**Fig. S1.**
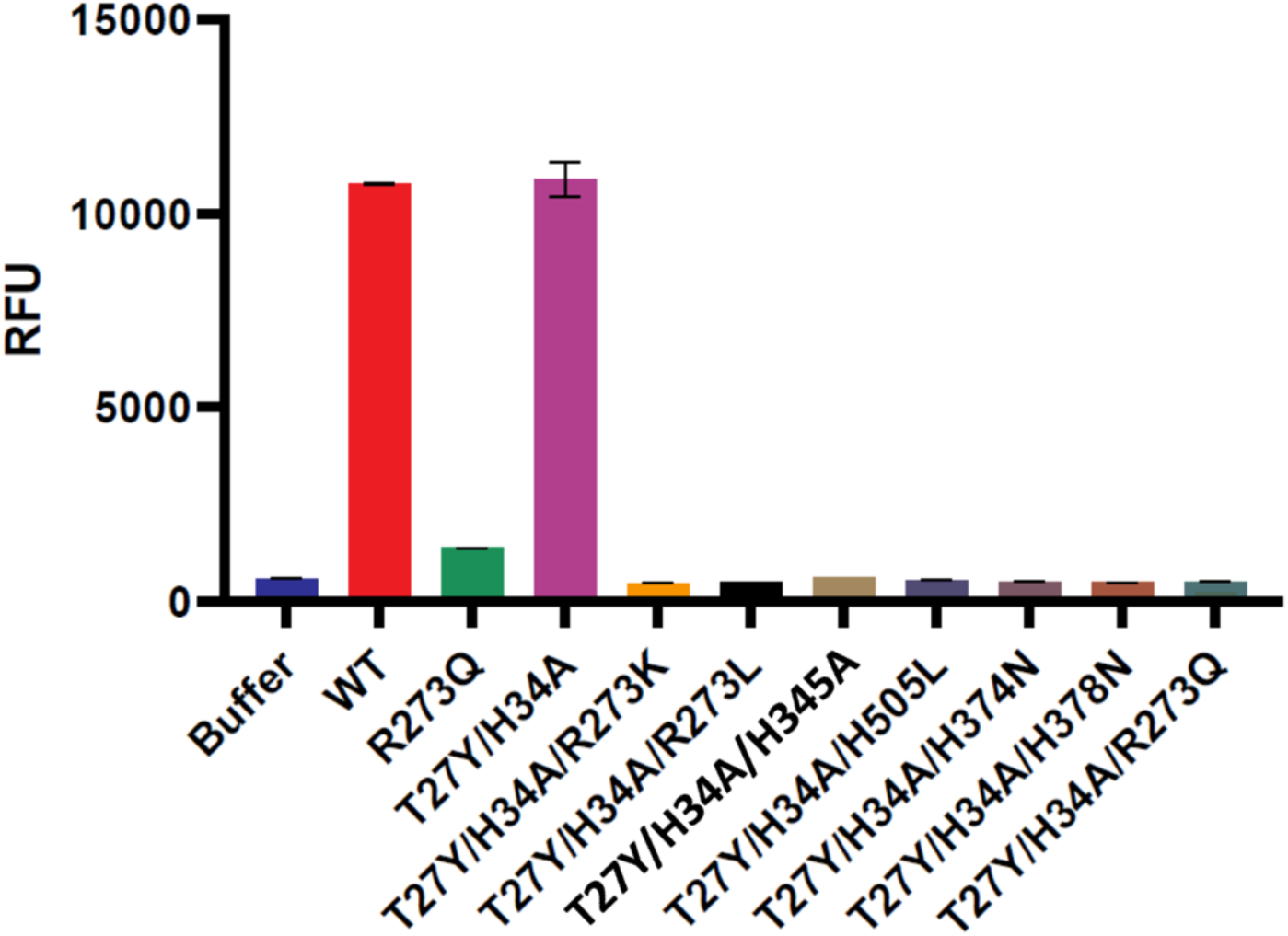
ACE2 activity assay. Enzyme activity of ACE2 in relative fluorescent units (RFU) for each decoy is shown.

To choose which of these activity-inhibiting mutations would be used in combination with the two affinity-enhancing mutations, we compared BLI kinetic analysis of S RBD binding for each. Of the triple mutants, the ones expressing R273K or H374N had the lowest dissociation coefficient (KD), that is, highest affinity binding.

The T27Y/H34A/H374N triple mutant was chosen for further testing because it showed better biophysical properties, including a lower propensity to aggregate, higher titer/better Tm as compared to the decoy with the R273K substitution.

**Table S1.** BLI analysis of S RBD binding by triple mutants. Titer analysis for the triple decoys is shown below in Fig. S2 and Table S2.

| Loading Sample ID | kon (1/Ms) | koff (1/s) | KD (nM) |
| --- | --- | --- | --- |
| T27Y/H34A | 3.77E+05 | 2.11E-04 | 0.56 |
| T27Y/H34A/R273K | 4.04E+05 | 1.79E-04 | 0.44 |
| T27Y/H34A/R273L | 4.06E+05 | 4.04E-04 | 1.00 |
| T27Y/H34A/H345A | 4.15E+05 | 3.95E-04 | 0.95 |
| T27Y/H34A/H505L | 4.09E+05 | 2.91E-04 | 0.71 |
| T27Y/H34A/H374N | 3.88E+05 | 2.44E-04 | 0.63 |
| T27Y/H34A/H378N | 3.95E+05 | 3.32E-04 | 0.84 |
| T27Y/H34A/R273Q | 4.86E+05 | 4.09E-04 | 0.84 |

**Fig. S2.**
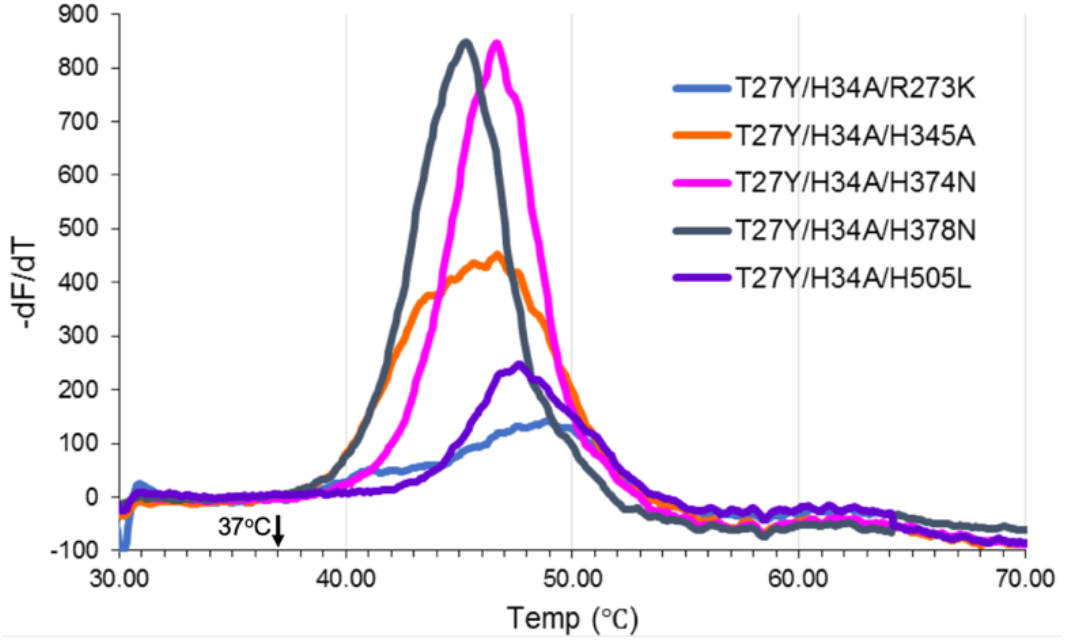
T_m_ analysis

**Table S2.** Tm analysis. The BLI kinetics analysis and binding values for ACE2 WT binding to naturally-occurring B.1.351 and CAL.20C variants are shown in Fig. S3 and Table S2.

| ACE2 mutants | Day 7 Titer (µg/mL) | % main peak post-ProA purification |
| --- | --- | --- |
| T27Y/H34A/R273K | 10.6 | 55.7 |
| T27Y/H34A/H345A | 27.3 | 70.8 |
| T27Y/H34A/H374N | 23.2 | 82.7 |
| T27Y/H34A/H378N | 30.1 | 75.8 |
| T27Y/H34A/H505A | 5.3 | 70.0 |

**Fig. S3.**
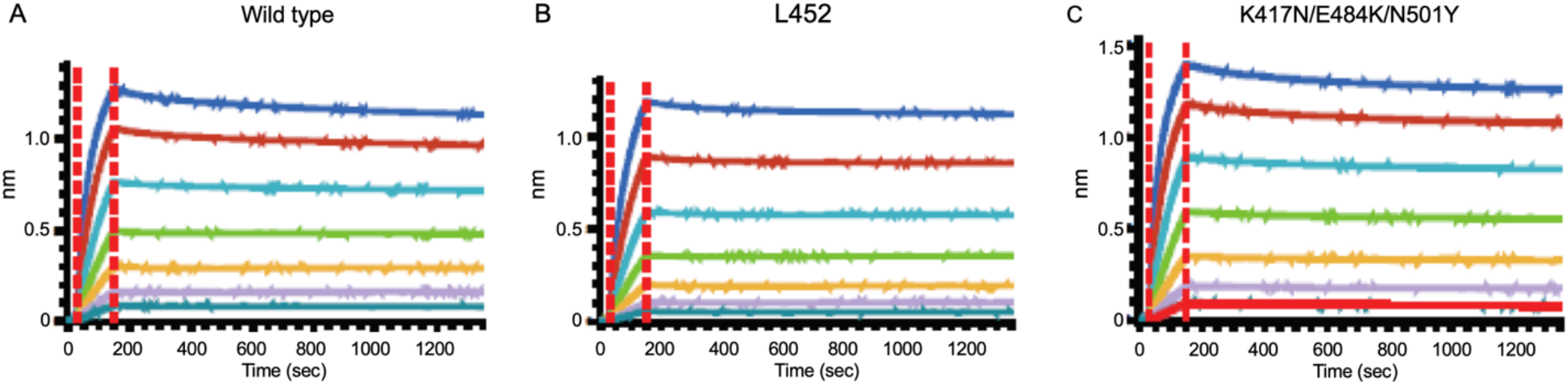
BLI kinetic analysis of ACE wild type (WT) decoy against SARS-CoV-2 RBD WT and L452R and K417N/E484K/N501Y mutants with avidity.

**Table S3.**
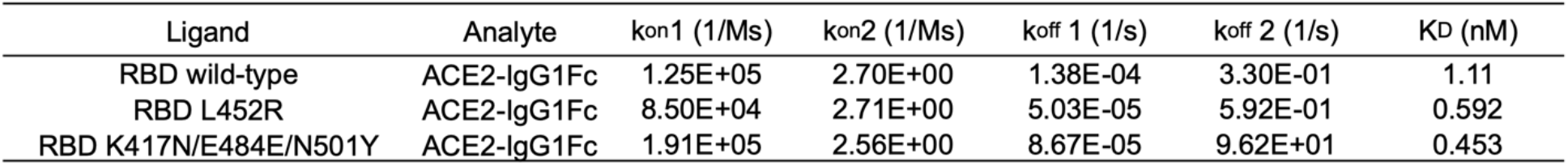
BLI kinetic analysis of ACE(WT) decoy against SARS-CoV-2 RBD mutants with avidity.

## Notes

### Competing Interest Statement

Authors listed with ImmunityBio, Inc. assisted with the creation or analysis of the ACE2 Triple Decoy (a potential therapeutic) described.

